# Erythropoietin alleviates intellectual disability and autism-like behavior of mice caused by *Zbtb20* haploinsufficiency, a construct-valid model of Primrose syndrome

**DOI:** 10.1101/2025.09.11.675680

**Authors:** Martin Hindermann, Justus BH Wilke, Yasmina Curto, Stefan N. Oline, Vinicius Daguano Gastaldi, Umer Javed Butt, Rakshit Dadarwal, Umut Çakır, Anja Ronnenberg, Kurt Hammerschmidt, Susann Boretius, Anastassia Stoykova, Anton B. Tonchev, Klaus-Armin Nave, Manvendra Singh, Hannelore Ehrenreich

## Abstract

Among the known genetic causes of syndromic autism spectrum disorders (ASD) are transcription factor deficiencies. In this regard, haploinsufficiency of the *‘zinc finger and broad complex, tramtrack, bric and brac domain-containing protein 20’ (ZBTB20)* leads to a prototypical clinical picture, referred to as Primrose syndrome, comprising severe ASD symptoms together with intellectual disability. Here, we present a comprehensive behavioral and phenotypical characterization of *Zbtb20^+/-^* mice, a construct valid model of this thus far untreatable human condition. *Zbtb20^+/-^* mice exhibit diminished sociability, reduced vocalization, distinct repetitive behaviors, impaired cognitive flexibility, hyperactivity and hypoalgesia. Magnetic resonance imaging reveals increased volumes of hippocampus, cerebellum, brain matter, and whole brain, confirmed by postmortem brain weight measurements. Due to our previous observation of enhanced ZBTB20 expression in CA1 pyramidal neurons upon recombinant human erythropoietin (rhEPO) injections, we anticipated a mitigating effect through rhEPO treatment of *Zbtb20* deficiency/Primrose syndrome. Indeed, after three weeks of alternate-day rhEPO injections, a remarkable improvement in the behavioral phenotype was observed. Our results highlight rhEPO as a first promising treatment for Primrose syndrome.

## Introduction

Autism spectrum disorder (ASD), with a prevalence of ∼1% across cultures, is characterized by 3 primary behavioral domains: Impaired sociability, reduced communication, and repetitive behaviors (Zoghbi & Bear, 2012). Additional symptoms are decreased cognitive flexibility (Lage et al., 2024), hyperactivity (Lai et al., 2019; Lugo-Marin et al., 2019), and pain hyposensitivity (Allely, 2013; Tordjman et al., 2009). Despite multifactorial causes, including dysfunctions of chromatin and transcription factors, the final common pathway of ASD converges at the synapse (De Rubeis et al., 2014). A transcription factor gene, associated with ASD is the *‘zinc finger and broad complex, tramtrack, bric and brac domain-containing protein 20’ (ZBTB20)*, located at chromosome 3q13.31. The encoded protein is particularly abundant in the hippocampus (Nielsen et al., 2014). *ZBTB20* contains 5 C2H2 zinc finger domains (ZnFI-ZnFV) and an N-terminal BTB (broad complex, tramtrack, bric and brac) domain, which facilitates interaction with the DNA (Sutherland et al., 2009).

ZBTB20 plays a crucial role in glucose metabolism, postnatal growth, neurogenesis (Sutherland et al., 2009; Xie et al., 2010; Zhang et al., 2012), and the specification of the medial pallium, which forms the hippocampus (Nielsen et al., 2010; Nielsen et al., 2007; Nielsen et al., 2014; Rosenthal et al., 2012; Xie et al., 2010). Furthermore, ZBTB20 is instrumental for maturation of hippocampal cornu ammonis 1 (CA1) neurons, the development of dendritic and synaptic structures, olfactory bulb neurogenesis, and the generation of neuronal layers in the developing cortex (Doeppner et al., 2019; Medeiros de Araujo et al., 2021; Ren et al., 2012; Tonchev et al., 2016).

In humans, a heterozygous pathogenic variant of *ZBTB20* causes the extremely rare, autosomal dominant Primrose syndrome (Primrose, 1982; Cordeddu et al., 2014) which is characterized by macrocephaly with developmental delay, intellectual disability, behavioral abnormalities, typical facial phenotypes, altered glucose metabolism, hypotonia, agenesis of the corpus callosum, hearing loss, ocular anomalies, cryptorchidism and calcification of the ear cartilage (Alby et al., 2018; Arora et al., 2020; Carvalho & Speck-Martins, 2011; Casertano et al., 2017; Cleaver et al., 2019; Cordeddu et al., 2014; Ferreira et al., 2019; Grimsdottir et al., 2019; Juven et al., 2020; Mathijssen et al., 2006; Mattioli et al., 2016; Melis et al., 2020; Yamamoto-Shimojima et al., 2020). Behavioral deviations include attention-deficit/hyperactivity disorder (ADHD), self-injurious behavior, sleep disturbances, tics, stereotypies, and ASD (Dalal et al., 2010; Melis et al., 2020; Rasmussen et al., 2014; Stellacci et al., 2018).

Regarding animal models, only few studies reported on a limited number of behavioral tests performed with *Zbtb20^+/-^* mice. These tests examined visual capability, hippocampus-dependent learning and memory, anxiety, exploration, nociception and circadian rhythm (Gulbranson et al., 2021; Jia et al., 2021; Nielsen et al., 2007; Qu et al., 2016; Ren et al., 2012). Thus far, however, a comprehensive behavioral characterization or any kind of treatment approach are completely lacking.

The glycoprotein erythropoietin (EPO), named after its initially discovered effects on the hematopoietic system, has over the last decades attracted attention due to its neuroprotective, neuroregenerative and cognition-enhancing properties. Moreover, EPO was just found to be a potent driver of neurodifferentiation (Singh et al., 2023). Recently, we detected an increase in ZBTB20 expression within the hippocampal CA1 region in mice following recombinant human EPO (rhEPO) treatment (Fernandez Garcia-Agudo et al., 2021).

We thus hypothesized that rhEPO application might augment the residual expression of ZBTB20 in heterozygous mice, and thereby mitigate the overall phenotype. To test this hypothesis, we conducted an in-depth behavioral and phenotypical characterization of *Zbtb20^+/-^* mice following 3 weeks of rhEPO application. The results revealed multifaceted improvements, suggesting rhEPO as a first promising treatment approach for Primrose syndrome.

## Methods

### Mice

All animal experiments were carried out with the approval of the local Animal Care and Use Committee (Niedersächsisches Landesamt für Verbraucherschutz und Lebensmittelsicherheit, LAVES, Oldenburg, Germany, license 42502-04-18/2803). **Both genders** were tested in this study. Unless otherwise specified, mice were separated by sex, genotype and treatment and housed in standard type II cages (Tecniplast, Hohenpleißberg, Germany) in groups of 3-5 within ventilated cabinets (Scantainers, Scanbur Karlsunde, Denmark) in a temperature- and humidity-controlled environment (∼22°C, ∼50%) with a 12 hour light-dark cycle (lights on at 7am) and water and food (Sniff Spezialdiäten, Bad Sodenberg, Germany) *ad libitum*. Cages were provided with wood-chip bedding and nesting material (Sizzle Nest, Datesand, Bredbury, United Kingdom). Group sizes were predicated on previous experience and under consideration of the RRR principle. All experiments were conducted by investigators unaware of group assignment (′fully blinded′).

The 129/Sv mice, carrying the *Zbtb20* mutant allele, have been previously described (Rosenthal et al., 2012). In this study, mice were bred by crossing C57BL/6N *Zbtb20^+/-^* males with C57BL/6N WT females. This ensured the birth of only *Zbtb20^+/-^* and *Zbtb20^+/+^* mice as littermates (control), thereby preventing the birth of neonatally lethal *Zbtb20^-/-^* animals. Mutations of the *Zbtb20* allele were confirmed by PCR-based genotyping (protocols available upon request). Mice were weaned at PND21 and separated by sex and genotype to avoid inclusion effects or aggressive behavior against potentially affected animals. Experiments started with the first cohort at an age of 4 weeks (PND28; males), and with the second cohort at an age of 7 weeks (PND50; females) and stretched over a period of ∼5 months each. Later, a (partial) replication and extension cohort was necessary, with females starting on PND28 and males on PND36.

### EPO/Placebo Injections

Animals were categorized into 2 groups based on sex, and further subdivided into 2 subgroups based on genotype. At PND28, 50% of the mice in each subgroup received a total of 11 intraperitoneal injections of 5000IU/kg rhEPO (NeoRecormon, Roche) or placebo (solvent solution, volume: 0.01ml/gBW), every other day over a period of 3 weeks (Adamcio et al., 2008).

### Behavioral Characterization

Behavioral experiments were performed during the light phase (unless stated otherwise) as reported in detail previously (Dere et al., 2014; El-Kordi et al., 2013; Hammerschmidt et al., 2012) and in the following order:

#### First (male) and second (female) cohort

Open Field, Hole Board, Y-Maze, Social Interaction in Pairs, 3-Chamber Sociability Test, Marble Burying Test, LABORAS and Nestbuilding Test, Vocalization Test, Prepulse Inhibition, Morris Water Maze, Forced Swim Test, and 4h Complex Wheel Running.

#### Replication and extension cohort (males; females)

Neophobia, Rotarod, Grip Strength, Marble Burying, Sucrose Preference Test, Buried Food Test, Tail Suspension Test, Hot Plate Test. Individual tests are described in detail in Supplementary Methods.

#### Thermography

Methodology including data extraction and processing have been described in detail previously (Seidel et al., 2020). Briefly, during Y-Maze, Social Interaction in Pairs, and 3-Chamber Sociability Test, an A655sc infrared thermography camera (FLIR Systems, Oregon, USA) was positioned above the respective arenas to capture images at a resolution of 640x480 pixels and frame rates of 5Hz (for Y-Maze and Sociability Test) or 25Hz (for the Social Interaction in Pairs), utilizing the ResearchIR software (FLIR Systems). OpenCV 4 in Python 3.6 was employed to extract thermal data. Images were loaded and normalized to values ranging from 0 to 255, with higher values indicative of higher temperatures. For each of the 3 tests, regions of interest (ROIs) were delineated for the extraction of thermal data. A binary mask, encompassing the whole body (including tail) was generated by applying intensity thresholding and processing steps to mitigate image noise. Consequently, a large cluster of interconnected pixels within these ROIs could be identified, forming the contour of the mouse. Owing to variations in shape and temperature, the whole-body area could subsequently be segmented into a central body region and a distinct tail region, thereby facilitating the extraction of the mean temperatures of both areas. The parameter assessed was the temporal variation in temperature, expressed as Centralization Index (CI), i.e. ratio of body temperature to tail temperature (Seidel et al., 2020).

#### Perfusion

Following the last test, i.e. Complex Wheel Running for 4 hours, mice were immediately anesthetized using Avertin (Sigma-Aldrich, Germany, 600mg/kg BW intraperitoneal). Subsequently, they were subjected to transcardial perfusion with Ringer solution for 5 min (Ringer Fresenius, Fresenius KABI, Bad Homburg, Germany), followed by 4% formaldehyde/PBS for 10 min.

### Multi-animal pose tracking

To analyze mouse behavior in the social interaction in pairs paradigm, we applied SLEAP 1.2.9 (Social LEAP, open source, (Pereira et al., 2022)) for pose and position tracking. A 15-node skeleton was used to label 1300 frames across all videos recorded for this experiment evenly divided between male and female subjects. This training package was used to train a top-down model with 2 components: A centroid model to locate the mice, and a centered instance model to identify individual nodes within a 384 pixel-wide cropped box. Those models were then used to run inference on all videos, and the output was manually proofread for switches. The resulting proofread inference was exported as h5 files for further analysis.

### Magnetic Resonance Imaging (MRI)

Mice were weighed, anesthetized with ketamine (75mg/kg BW) and medetomidine (1mg/kg BW), followed by endotracheal intubation. A specialized respirator, specifically designed for rodents (Animal Respirator Advanced, TSE Systems, Bad Homburg, Germany), was employed for controlled ventilation, supplying room air, enriched with oxygen and mixed with isoflurane (0.5-2%). A pressure sensor allowed monitoring breathing rate, potential spontaneous breathing, or any movements. Body temperature was controlled using a rectal thermometer and regulated with a warm water circulation system. Mice were positioned in the MRI with heads secured in a tooth and palate holder (Boretius et al., 2009). MRI was performed at a magnetic field strength of 9.4 Tesla using a 4-channel receive-only mouse head coil (Biospec, Bruker BioSpin, Ettlingen, Germany). For the purpose of volumetric analyses, magnetization transfer (MT)-weighted images were acquired with a 3D fast low-angle shot (FLASH) sequence [with Echo Time/Repetition Time (TE/TR) = 3.4/15.2ms, flip angle 5°, Gaussian-shaped off-resonance pulse (off-resonance frequency 7.5ppm, RF power 6µT), and 100µm isotropic spatial resolution].

#### MRI Data Analyses

For volumetric analyses, MT-weighted images were converted to the NIfTI format, then preprocessed using denoising and bias field correction techniques (Avants et al., 2011). Preprocessing aimed at generating an unbiased anatomical population template. This was achieved using the optimised ANTs template construction pipeline (GitHub, CoBrALab). To visualize volumetric changes, nonlinear deformation fields of extracted brain images were utilized to create voxel-wise Jacobian determinants. To quantify the volume of specific brain areas, regions of interest (ROIs) were identified, including olfactory bulb, hippocampus, thalamus, corpus callosum, ventricles, cerebellum, and whole brain. These ROIs were determined on the study template through a process of manual segmentation, facilitated by software package AMIRA (Visage Imaging GmbH, Berlin, Germany). Subsequently, these ROIs were retransformed into the subject space. Each of these retransformed ROIs was individually checked and manually corrected if necessary. Finally, volume information was extracted from these corrected ROIs and used for statistical analyses and illustration purposes.

#### Organ Collection

Immediately following MRI, mice were weighed - still under anesthesia - and sacrificed for organ collection, including brain, heart, and lungs. Organs were weighed and stored in 2ml Eppendorfs until subjected to lyophilization, carried out at -56°C for 48 h under a vacuum pressure of 0.2 mBar, utilizing the Christ LMC-1 BETA 1-16 lyophilizer. In addition, tibiae were taken as secondary reference point (after BW) to create weight and size-adjusted ratios for the various organs.

### Immunohistochemistry

Mice were deeply anesthetized with Avertin (Sigma-Aldrich, Germany, 600mg/kg BW intraperitoneal) and transcardially perfused via left cardiac ventricle with Ringer’s solution followed by 4% PFA in PBS (0.1M, pH 7.4). Dissected brains were postfixed overnight in 4% PFA and frozen at -80°C after cryoprotection with 30% sucrose. Coronal sections of 30μm thickness were cut with cryostat Leica CM1950 (Leica Microsystems) and stored at -20°C in cryoprotective solution (25% ethylene glycol and 25% glycerol in PBS). For IHC, sections were permeabilized in PBS containing 0.3% Triton X-100 (PBST; Sigma-Aldrich) and blocked with 5% normal horse serum (NHS; Jackson ImmunoResearch Laboratories) for 1 hour at RT. Subsequently, slices were incubated with main primary antibody anti-ZBTB20 (rabbit, 1:500; Synaptic Systems 362003) diluted in 5% NHS with 0.3% PBST over 3 nights at 4°C. After washing, secondary antibody goat anti-rabbit Alexa 555 (A-21428; 1:500; Invitrogen) was incubated in 3% NHS with 0.3% PBST for 2 hours at RT. Sections were finally counterstained with 4′,6-diamidino-2-phenylindole (DAPI, 1:5000; Millipore-Sigma) and mounted on SuperFrostPlus Slides (Epredia) with Aqua-Poly/mount (Polysciences, Inc). A total of 12 mice (6 *Zbtb20* WT and 6 *Zbtb20^+/-^*) were used to assess the hippocampus. Images were obtained with a TCS-SP5 inverted system (Leica) equipped with a 20x objective (NA = 0.70). Representative images were taken at Bregma -1,58 mm.

### Quantitative RT-PCR

Hippocampal RNA was extracted from either cryosections (4 sections per mouse) or from whole hippocampus. The extraction was performed using the NucleoSpin totalRNA FFPE Kit (product number 740982, Macherey-Nagel, Duren, Germany) or the miRNeasy Mini Kit (QIAGEN, Hilden, Germany), respectively. The cDNA was then synthesized using the SuperScript III Reverse Transcriptase (Thermo Fisher Scientific Life Technologies GmbH, Darmstadt, Germany). This process involved the use of 1µg of RNA, along with oligo (dT) and Random Hexamer Primer, in a total reaction volume of 20µl. Subsequently, RT-qPCR was performed using 5µl of Power SYBR Green PCR Master Mix (product number 4367660, Thermo Fisher Scientific). This procedure utilized 4µl of 1:10 diluted cDNA as template and 1pmol of primers (dissolved in 1µl of H_2_O). Fold changes in gene expression were calculated with the ΔΔCt method, using *β-actin* and *Hprt1* as reference genes. The qPCR reactions (3 technical replicates) were run on LightCycler® 480 System (Roche, Mannheim, Germany).

*Zbtb20* forward primer: 5′-CAGCCCTCATCCACTCGAC-3′

*Zbtb20* reverse primer: 5′-TCCCCTTGCAACTGATGTCAC-3′

*β-actin* forward primer: 5′-CTTCCTCCCTGGAGAAGAGC-3′

*β-actin* reverse primer: 5′-ATGCCACAGGATTCCATACC-3′

*Hprt1* forward primer: 5′-GCTTGCTGGTGAAAAGGACCTCTCGAAG-3′

*Hprt1* reverse primer: 5′-CCCTGAAGTACTCATTATAGTCAAGGGCAT-3′

### High parameter flow cytometry

Frequency of 6 major and 19 minor immune cell subsets were quantified in blood, bone marrow, lymph node, and spleen samples of wildtype versus heterozygous *Zbtb20^+/-^* mice as described in the Supplementary Methods. Pairwise comparisons between *Zbtb20^+/-^* and wildtype mice were performed using either the Wilcoxon-rank sum test with continuity correction or unpaired 2-sided t-tests with or without Welch’s correction depending on data normality and variance. To account for multiple testing, p-values were adjusted within each organ for all cell populations using the Benjamini–Hochberg false discovery rate (FDR) procedure. Both raw p-values and FDR-adjusted p-values are reported. Multivariate analysis of the global immune cell composition was performed using principal component analysis (PCA) on z-scored features, with 95% confidence ellipses plotted per genotype and organ. Group differences in overall immune profiles were assessed separately within each organ by permutational multivariate analysis of variance (PERMANOVA) based on Euclidean distances of the scaled data.

### snRNA-seq

This study was commenced by fetching the normalized data available from our previous study (Singh et al., 2023). We reanalyzed the observed lineages to provide a holistic catalogue at the finer resolution. After the batch correction, we performed unsupervised clustering of the single-cell gene expression profiles of all hippocampal samples, which identified 19 clusters at a resolution of 0.4 (preset by Seurat tool). We then examined the top differentially expressed genes for each cluster relative to all other clusters which enabled us to calculate the expression and differential expression of *Zbtb20.* UMAPs, Feature plots, Violin plots and Heatmaps were constructed using default functions of tools we implemented, except that we set the color scale manually. The detailed code to regenerate all the figures and analysis corresponding to this snRNA-seq dataset is available at https://github.com/Manu-1512/Zbtb20/.

### Statistical Analyses

Unless specified otherwise, statistical analyses were conducted utilizing Prism9 software (GraphPad Software) and results are presented as mean values ± standard deviations (SD). The normality of data distribution was assessed using the Shapiro-Wilk test, with an alpha error threshold set at 0.05. Depending on the distribution of the data, either 2-tailed unpaired Welch’s corrected t-tests or Mann-Whitney U-tests were employed to facilitate comparisons between groups. In the case of repeated measure data, a mixed-model Analysis of Variance (ANOVA) was applied. Statistical significance was determined based on p values < 0.05.

## Results

### Synopsis

This study was designed with the objective of performing an exhaustive behavioral characterization of *Zbtb20^+/-^* mice, a construct-valid model of the thus far untreatable human Primrose syndrome. Based on our previous observations of EPO specifically targeting ZBTB20 expression, we incorporated a 3-week treatment trial using rhEPO. The findings unveiled impairments across all domains that are typically associated with ASD and intellectual disability. Importantly, improvements in several pathological parameters were observed following rhEPO administration (figures 1-6; supplementary figures 1-3; supplementary tables 1-4), thus potentially offering a first promising therapeutic approach to Primrose syndrome.

**Figure 1:**
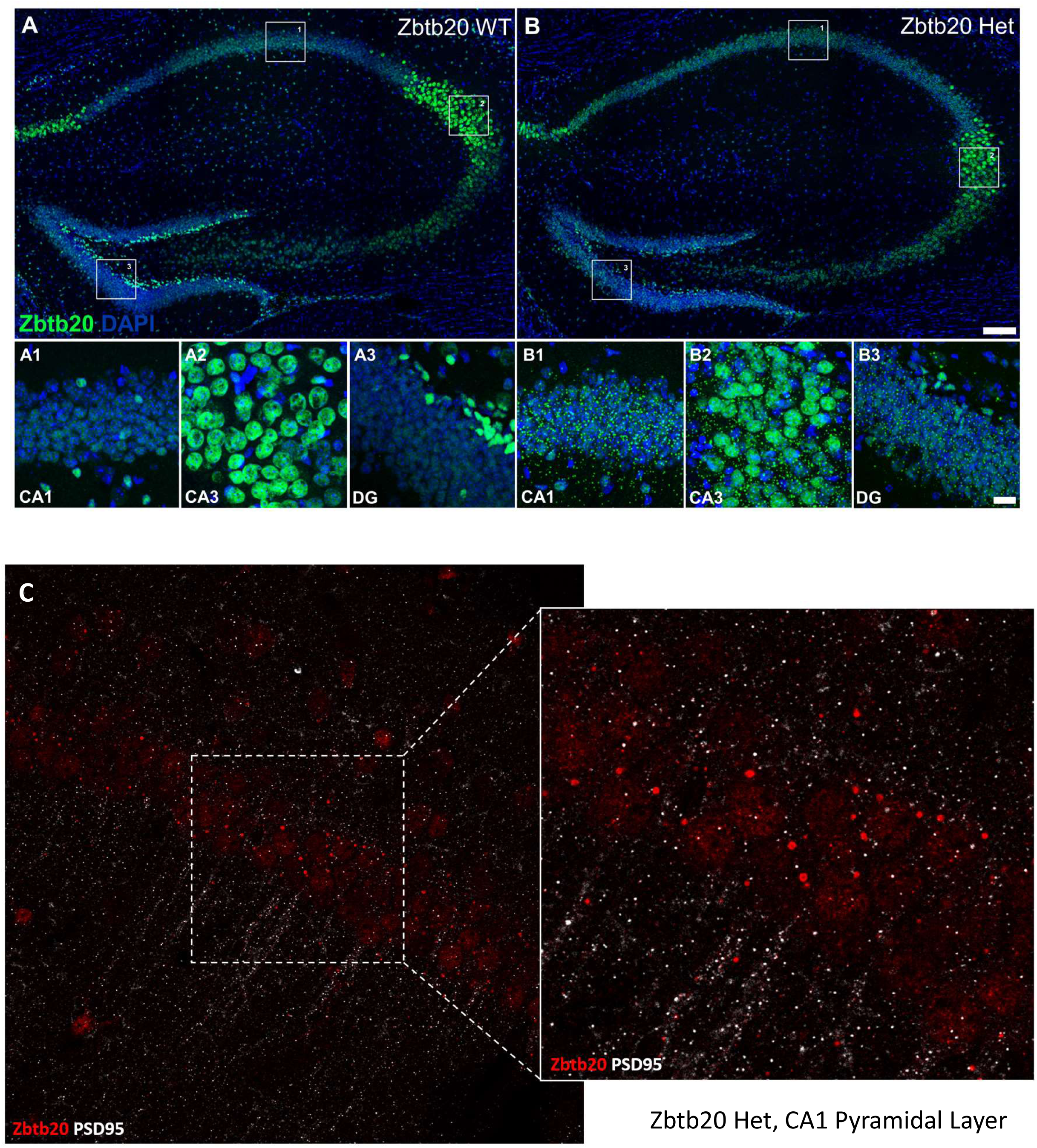
Histology. **(A-B):** Representative images of ZBTB20 expression (green) in the hippocampal region of *Zbtb20^+/+^* (WT) and *Zbtb20^+/-^* (Het) mice. **A1-A3** & **B1-B3**: Magnifications of the squared zones in **A** and **B**, representing CA1, CA3 and DG areas. Scale bar: 100µm for **A-B**; 20µm for magnifications. **(C):** CA1 Pyramidal Layer of *Zbtb20^+/-^* (Het) including magnification reveals no double-labeling of Zbtb20 with PSD95.

### Immunohistochemistry illustrates ZBTB20 expression in the hippocampus of *Zbtb20^+/+^* (WT) versus *Zbtb20^+/-^* (Het) mice

Immunohistochemical images of ZBTB20 expression in the hippocampal regions CA1, CA3 and DG of *Zbtb20^+/+^* (WT) and *Zbtb20^+/-^* (Het) mice document the expectedly reduced expression/green signal in Het mice (figure 1A-B). Interesting, but still unclear in nature, are the small green puncta distributed all over the hippocampus exclusively of Het mice. Exploratory co-staining for PSD95 excludes a simple co-localization of these puncta with synaptic structures (figure 1C). Furthermore, they are mainly located outside the nucleus, are not associated with astrocytic (GFAP) nor with neuronal processes (MAP2) (data not shown).

### Hematopoietic cell composition in lymphoid organs is largely unaffected in Zbtb20+/- mice

While ZBTB20 has been most extensively studied in the central nervous system, accumulating evidence, derived from cell-type specific *Zbtb20* knockout mice, highlights important roles for this transcription factor in several immunological processes (Chevrier et al., 2014; Krzyzanowska et al., 2022; Liu et al., 2013; Stoyanov et al., 2023). To assess whether ZBTB20 haploinsufficiency impacts the hematopoietic cell composition, we performed high-parameter flow cytometric immunophenotyping to quantify 6 major immune cell types, namely B cells, CD4+ and CD8+ T cells, NK cells, monocytes, and granulocytes, as well as 19 functionally distinct immune cell subsets across several lymphoid compartments, including blood, bone marrow, lymph nodes, and spleen (supplementary figure 3A and supplementary table 4). Qualitative comparison of immune cell clusters expectedly revealed major differences between the immune cell compartments but no gross abnormalities in *Zbtb20^+/-^* versus wildtype mice (supplementary figure 3B). Subsequent quantitative PERMANOVA comparison of the immune cell compositions within each lymphoid compartment confirmed the absence of major changes in the immune cell composition of *Zbtb20^+/-^* mice (all p>0.05, supplementary figure 3C). Detailed analysis of individual immune cell populations revealed subtle but significant increase in CD4+ T cells in lymph nodes of *Zbtb20^+/-^* mice (FDR=0.023, fold change = 1.1) as well as an increase in splenic NK cells (FDR = 0.0275, fold change = 1.2). No significant differences between *Zbtb20^+/-^* and wildtype mice were observed for distinct subsets of NK cells, T cells, B cells, and monocytes (all FDR>0.05, supplementary table 4).

### *Zbtb20^+/-^* males display impairments in all primary domains of ASD-like behavior

An *in-depth* behavioral characterization of *Zbtb20^+/-^* males (figure 2A-B) revealed impairments across all primary domains of an ASD phenotype. The compromised social interaction became evident in the 3-chamber sociability test. Here, placebo-treated *Zbtb20^+/-^* males failed to differentiate between the chamber housing a stimulus mouse and the opposite chamber containing an empty cage (p=0.1261). In contrast, both placebo and rhEPO-treated *Zbtb20^+/+^* and rhEPO-treated *Zbtb20^+/-^* demonstrated a preference for the chamber of the stimulus mouse over the empty chamber (p=0.0008, p<0.0001, p=0.0080, figure 2C and supplementary table 1).

**Figure 2:**
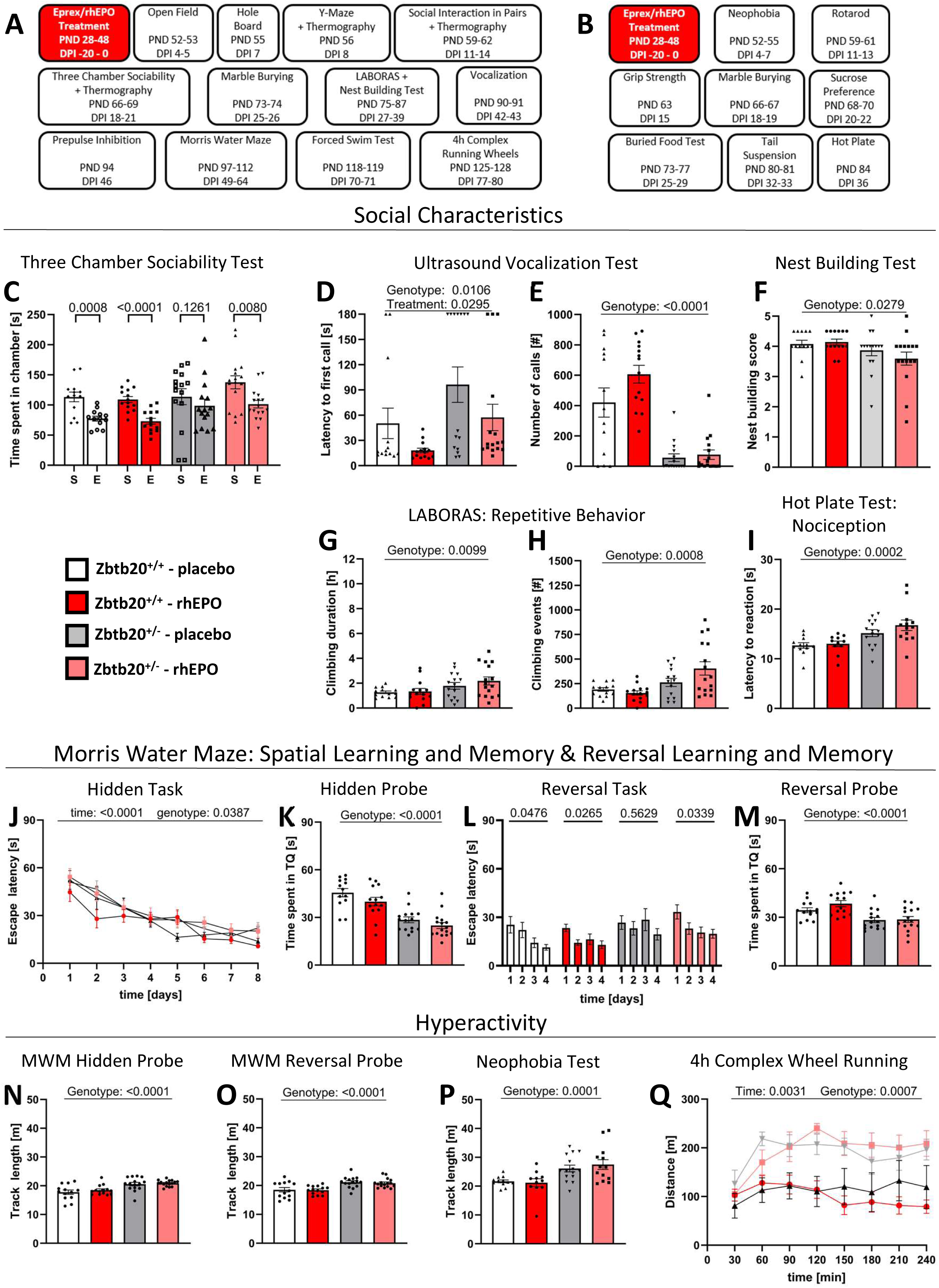
Experimental outline and *Zbtb20^+/-^* impairments in ASD domains. **(A-B)** Graphical experimental outline for main **A** and additional **B** cohort of male *Zbtb20^+/-^* mice and their *Zbtb20^+/+^* littermates, used as control. After 11 consecutive placebo resp. rhEPO injections, behavior experiments were performed (PND=postnatal day, DPI=days post injection). **(C-F)** Assessment of social characteristics shows impaired social preference in the 3-chamber sociability test **C**, since placebo-treated *Zbtb20^+/-^* mice did not distinguish between stimulus mouse (S) and empty cage (E). This impairment is rescued by rhEPO treatment (t-test, mean±SEM). Communication assessed with ultrasound vocalization **D-E** shows that *Zbtb20^+/-^* mice exhibit an increased latency to first call **D** with improvement after rhEPO treatment and a reduced number of calls **E** compared to controls (2-way ANOVA, mean±SEM). Nest building **F** results in a reduced score in *Zbtb20^+/-^* mice regarding quality of constructed nests overnight (2-way ANOVA, mean±SEM). **(G-H)** Repetitive behavior was assessed in home cages via LABORAS and shows increased duration **G** and events **H** of climbing in *Zbtb20^+/-^* mice (2-way ANOVA, mean±SEM). **(I)** Hot plate test reveals hyposensitivity towards heat-mediated nociception in *Zbtb20^+/-^* mice with an increased latency of the first reaction to heat perception (2-way ANOVA, mean±SEM). **(J-M)** Spatial learning, memory, as well as reversal learning and memory were assessed in Morris Water Maze. Hidden training over 8 days shows time and genotype effect in escape latency (**J,** 3-way ANOVA, mean±SEM). Probe trial for hidden learning task **K** reveals impaired spatial memory in *Zbtb20^+/-^* mice compared to controls, shown by reduced time spent in target quadrant (TQ, 2-way ANOVA, mean±SEM). Reversal hidden training over 4 days **L** displays impaired spatial reversal learning in placebo-treated *Zbtb20^+/-^* mice and a rescue effect after rhEPO treatment, indicated by escape latency to platform (ANOVA, mean±SEM). Reversal probe trial **M** demonstrates impaired reversal spatial memory in *Zbtb20^+/-^* mice, indicated by the time spent in target quadrant (2-way ANOVA, mean±SEM). **(N-Q)** Hyperactivity of *Zbtb20^+/-^* mice was assessed in probe trials for both hidden **N** and reversal **O** tasks of Morris Water Maze, neophobia test (**P,** all 2-way ANOVA, mean±SEM)) and 4h-complex wheel running (**Q**, 3-way ANOVA, mean±SEM).

Communication deficits were discernible in the vocalization test. Both placebo and rhEPO-treated *Zbtb20^+/-^* males exhibited an increased latency to their first vocalization upon exposure to an anaesthetized female stimulus mouse (genotype effect: p=0.0106). Interestingly, rhEPO-treatment of both *Zbtb20^+/+^* and *Zbtb20^+/-^* mice reduced the latency to their first vocalization (treatment effect: p=0.0295). Furthermore, both *Zbtb20^+/-^* groups emitted significantly fewer calls towards the stimulus mice throughout the 3-min recording, compared to their respective controls (genotype effect: p<0.0001, figure 2D-E, and supplementary table 1). In nest building, another social skill test, *Zbtb20^+/-^* males constructed nests of inferior quality compared to WT controls and rhEPO had no effect (genotype effect: p=0.0279, figure 2F and supplementary table 1).

This result was also mirrored in their female littermates, where both *Zbtb20^+/-^* groups scored lower in nest building compared to their respective WT controls (genotype effect: p<0.0001, supplementary figure 1A and supplementary table 2). Intriguingly, the diminished quality of nests constructed by both *Zbtb20^+/-^* genders was already obvious in their home cages, an observation that experimenters became aware of after being unblinded at project conclusion.

LABORAS exposure revealed enhanced repetitive behavior, with *Zbtb20^+/-^* males displaying increased duration of climbing and number of climbing events (genotype effects: p=0.0099 and p=0.0008, respectively; figure 2G-H and supplementary table 1) compared to WT controls. In females, the result was similar (genotype effects: p=0.0145 and p=0.0002, respectively; supplementary figure 1B-C and supplementary table 2).

### Additional abnormalities of *Zbtb20^+/-^* males linked to ASD-like behavior

In the hot plate test, an augmented tolerance for heat-mediated pain became aware. Both placebo and rhEPO-treated *Zbtb20^+/-^* males exhibited a heightened latency to their initial visible reaction towards the stimulus (genotype effect: p=0.0002, Fig2I and suppl. table 1). In females, placebo-treated *Zbtb20^+/-^* mice also demonstrated an increased latency compared to their WT control (genotype effect: p=0.0003). However, unlike in males, female rhEPO-treated *Zbtb20^+/-^* mice displayed a latency to the first reaction that was on par with WT controls and therefore lower than placebo-treated *Zbtb20^+/-^* females (treatment effect: p=0.0093, suppl. Fig 1D and suppl. table 2).

The Morris Water Maze was employed to evaluate spatial learning and memory. *Zbtb20^+/-^* males exhibited visual abilities equivalent to their WT controls (suppl. table 1). A genotype effect was observed during 8 days of hidden training (p=0.0387, Fig 2J). During both hidden and reversal probe trials, *Zbtb20^+/-^* males spent significantly less time in the target quadrant that previously held the escape platform (both genotype effects: p<0.0001, Fig 2K-M and suppl. table 1). Between both probe trials, 4 days of reversal training were conducted, revealing that placebo-treated *Zbtb20^+/-^* males were the only group not showing significant improvement in escape latency over time (p=0.5629). In contrast, both placebo and rhEPO-treated *Zbtb20^+/+^* males (p=0.0476, p=0.0265) and rhEPO-treated *Zbtb20^+/-^* males improved significantly (p=0.0339, Fig 2L and suppl. table 1). In females, a phenotype was only observable during both probe trials of hidden and reversal task where *Zbtb20^+/-^* mice spent less time in the target quadrant compared to controls, independent of treatment (genotype effects: p=0.0153, resp. p=0.0040, suppl. Fig 1F-H and suppl. table 2). Both male and female *Zbtb20^+/-^* mice displayed increased locomotion across a variety of tests. In males, hidden and reversal probe trials (genotype effects: p<0.0001), the neophobia test (genotype effect: p=0.0001), and 4-hour complex wheel running (genotype effect: p=0.0007, Fig 2N-Q and suppl. table 1) revealed significantly increased track lengths. In females, *Zbtb20^+/-^* mice demonstrated increased locomotion in the open field test (genotype effect: p=0.0006), both hidden and reversal probe trials (genotype effects: p=0.0009, resp. p=0.0004), and in the LABORAS test (genotype effect: p=0.0076, suppl. Fig 1L-O and suppl. table 2).

### *Zbtb20^+/-^* phenotypes in working memory, sensorimotor gating, and motivation

The Y-Maze test was employed to assess working memory by contrasting the frequency of alternating and non-alternating entries into the maze arms. Despite all 4 male groups performing significantly more alternating than non-alternating entries, a comparison of the deltas from these 2 readouts revealed that placebo-treated *Zbtb20^+/-^* mice had the lowest delta of all groups, significantly lower than rhEPO-treated *Zbtb20^+/-^* males, thus demonstrating a treatment effect (p=0.0314, Fig 3A-B and suppl. table 1). The open field and hole board tests were utilized to assess exploratory behavior and the motivation to engage in such. *Zbtb20^+/-^* males spent more time within the periphery and less time within the intermediate zone in the open field compared to their WT controls (genotype effects: p=0.0085, resp. p=0.0040, Fig 3C and suppl. table 1). The hole board experiment revealed a reduction in revisits to the most recently visited hole in placebo-treated *Zbtb20^+/-^* males compared to their WT controls (genotype effect: p=0.0148). Additionally, an increase towards control levels was observed in rhEPO-treated *Zbtb20^+/-^* (treatment effect: 0.0139, Fig 3D and suppl. table 1). A similar result was obtained in the female groups, where placebo-treated *Zbtb20^+/-^* mice showed fewer revisits compared to WT control, and rhEPO-treated *Zbtb20^+/-^* females exhibited a tendency of increasing the number of revisits to control level (genotype effect: p=0.0982, treatment effect: 0.0526, suppl. Fig1I and suppl. table 2).

**Figure 3:**
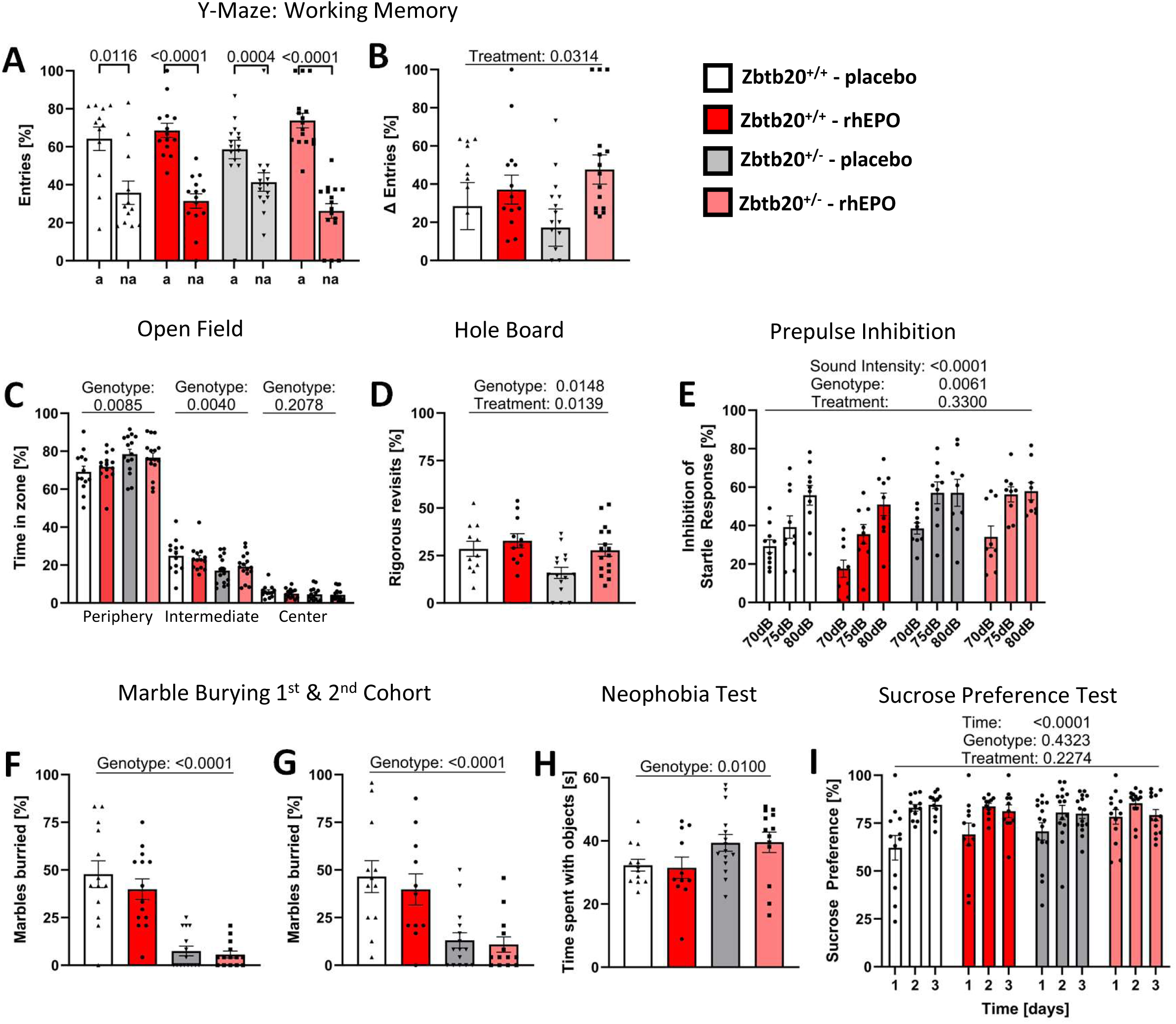
Working memory, exploration, PPI, obsessive-compulsive readouts and related control testing in *Zbtb20^+/-^*mice. **(A-B)** Working memory was assessed in the Y-Maze test and shows significantly more alternating (a) than non-alternating (na) entries in all 4 groups (**A,** t-test, mean±SEM). Comparing delta of entries **B** reveals further improved preference of alternating entries in rhEPO-treated *Zbtb20^+/-^* mice compared to placebo-treated controls (2-way ANOVA, mean±SEM). **(C)** Placebo-treated *Zbtb20^+/-^* mice show increased time spent in peripheral zone and reduced time spent in intermediate zone in the open field test (ANOVA, mean±SEM). **(D)** Explorative behavior was also assessed in hole board test, revealing reduced rigorous revisits of placebo-treated *Zbtb20^+/-^* mice and a rescue effect upon rhEPO treatment (2-way ANOVA, mean±SEM). **(E)** Prepulse inhibition test shows a genotype effect, namely increased inhibition of startle response in both placebo and rhEPO-treated *Zbtb20^+/-^* mice (3-way ANOVA, mean±SEM). **(F-G)** Marble burying test resulted in reduced numbers of buried marbles in the main **F** and, as control for this unexpected result, in an additional **G** cohort (2-way ANOVA, mean±SEM). **(H)** Neophobia test shows increased interaction time with objects in placebo-treated *Zbtb20^+/-^* mice and no treatment effect (2-way ANOVA, mean±SEM). **(I)** Sucrose preference test demonstrates absence of anhedonia-like (depressive) behavior in placebo and rhEPO-treated *Zbtb20^+/-^* mice (3-way ANOVA, mean±SEM).

In the prepulse inhibition paradigm, which was used to evaluate sensorimotor gating, significant distinctions were observed among the various sound intensities in all 4 groups (sound intensity: p<0.0001). Moreover, a genotype effect was identified when comparing both placebo and rhEPO-treated *Zbtb20^+/-^* males with their respective WT controls (p=0.0061, Fig 3E and suppl. table 1). Interestingly, in the marble burying test, both cohorts of placebo and rhEPO-treated *Zbtb20^+/-^* males buried fewer marbles compared to their respective WT controls (both genotype effects: p<0.0001, Fig 3F-G and suppl. table 1). A similar result was observed in females, where placebo and rhEPO-treated *Zbtb20^+/-^* mice of both cohorts buried significantly fewer marbles compared to their respective controls (genotype effects: p<0.0001, resp. p=0.0034, suppl. Fig 1J-K and suppl. table 2). To further investigate potential causes for these unusual results, additional tests for the assessment of neophobia, depression-like behavior, and anhedonia were conducted in a second experimental cohort of this project. In neophobia testing, both male and female *Zbtb20^+/-^* mice did not show decreased time spent with objects compared to their controls (suppl. tables 1&2), but *Zbtb20^+/-^* males exhibited an increased time spent with novel objects (genotype effect: p=0.0100, Fig 3H and suppl. table 1). For depression-like behavior, tail suspension and forced swim tests were performed but did not reveal an overall increased time of immobility in *Zbtb20^+/-^* males nor females (suppl. tables 1&2). For the assessment of anhedonia, the sucrose preference test was conducted, showing all 4 groups with significantly increasing preference towards sucrose over the experimental time of 3 days (time effect: p<0.0001, Fig 3I and suppl. table 1). Females also revealed no differences across groups in this test (suppl. table 2).

### MRI uncovered structural differences for brain regions

A voxel-wise Jacobian determinants map was created to provide a visual representation of volumetric alterations in the mouse brain (Fig 4A-B). This map intensely delineates the increasing (yellow/red) and decreasing (blue/white) volumes when comparing both placebo-treated groups (Fig 4A) and both placebo and rhEPO-treated *Zbtb20^+/-^* groups (Fig 4B). This comprehensive overview, combined with findings reported from previous studies and the technical prowess of this MRI technique, facilitated the volume analysis of an array of brain regions.

**Figure 4:**
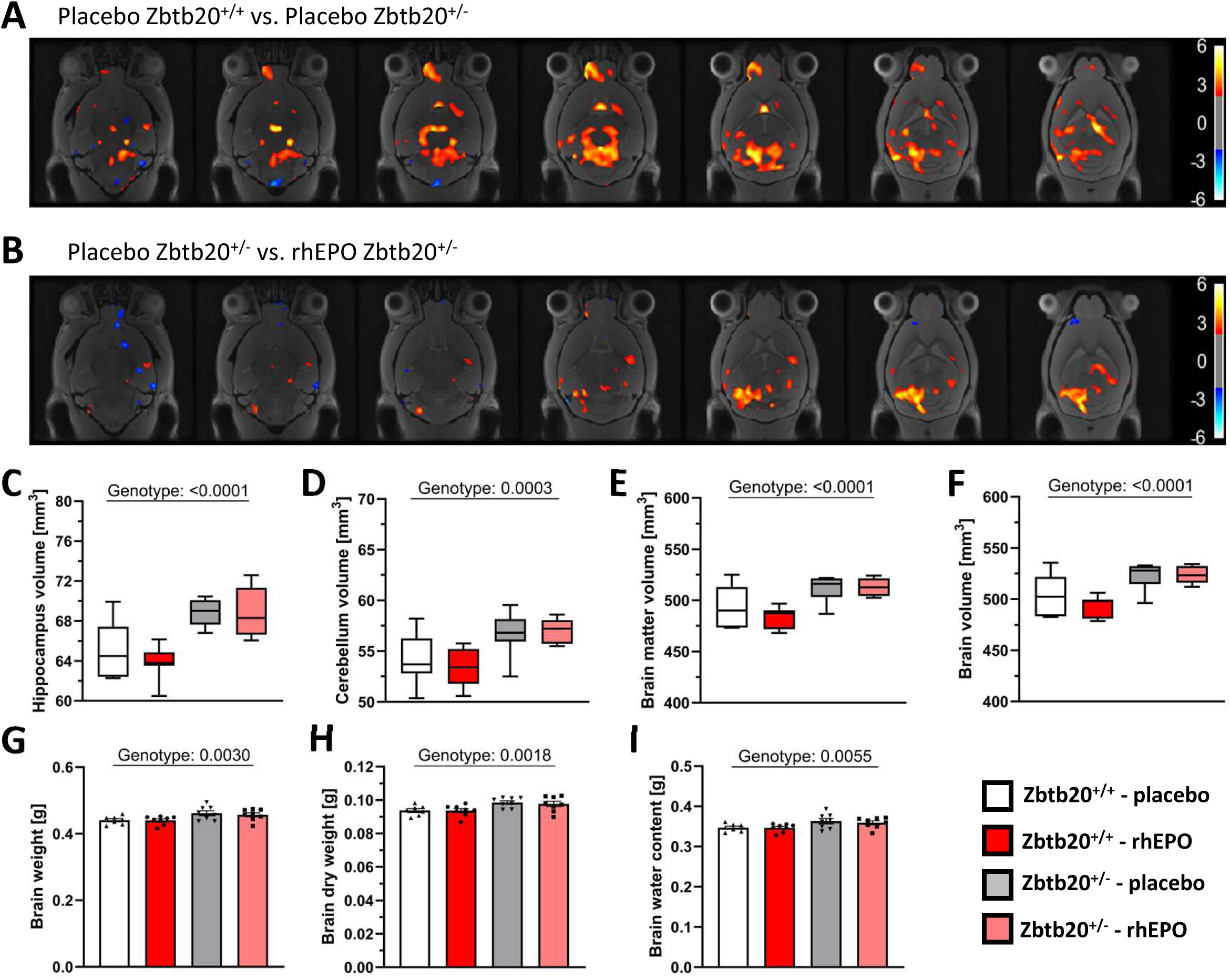
Magnetic resonance imaging (MRI) **(A-B)** Voxel-wise Jacobian determinants map indicates increased (red/yellow) or decreased (blue) volume changes in both placebo-treated *Zbtb20^+/+^* vs *Zbtb20^+/-^* **A** and placebo- vs rhEPO-treated *Zbtb20^+/-^* **B** mice. **(C-F)** Volumetric analyses of specific brain regions show in both placebo- vs rhEPO-treated *Zbtb20^+/-^* mice increased volume of hippocampus **C**, cerebellum **D**, brain matter **E**, and whole brain **F**. **(G-I)** Weight assessment of whole brains. **G** Immediately after extraction, brains of both placebo and rhEPO-treated *Zbtb20^+/-^* mice display increased weight compared to controls. **H** Likewise, after 48 hours of lyophilization, brain dry weight of both placebo and rhEPO-treated *Zbtb20^+/-^* mice is enhanced. **I** Also, calculated water content is augmented in both placebo and rhEPO-treated *Zbtb20^+/-^* mice (2-way ANOVA; mean±SEM).

In males, both *Zbtb20^+/-^* groups exhibited an increased volume of hippocampus (p<0.0001), cerebellum (p=0.0003), brain matter (p<0.0001), and whole brain (p<0.0001, Fig 3C-F and suppl. table 1). The brain weight immediately post-extraction, dry weight, and computed water content corroborated these findings (genotype effects: p=0.0030, p=0.0018, p=0.0055, Fig 4G-I and suppl. table 1). The brain regions of the olfactory bulb, thalamus, corpus callosum, and ventricles unveiled no volumetric differences (suppl. table 1).

The volumetric analysis in females disclosed a diminished olfactory bulb volume in both placebo and rhEPO-treated *Zbtb20^+/-^* mice (genotype effect: p<0.0001, suppl. Fig2A and suppl. table 2). While the volumes of both hippocampus and cerebellum were increased in placebo-treated *Zbtb20^+/-^* mice (genotype effect: p=0.0060, resp. p=0.0047), rhEPO treatment led to a reduction of the volume (treatment effects: p=0.0293, resp. p=0.0156, suppl. Fig2B-D and suppl. table 2). Volumetric changes of the ventricles revealed a treatment-dependent reduction (treatment effect: p=0.0007, suppl. Fig2C and suppl. table 2).

### snRNA-seq discovered upregulation of ZBTB20 in pyramidal neurons on rhEPO

To begin examining the ZBTB20 expression in the mouse hippocampus, we leveraged the recently published single nuclei RNA-seq (snRNA-seq) dataset encompassing a broad range of neuronal and non-neuronal cell types (Singh et al., 2023). By refining these lineages deeper using the preset resolution of 0.4 from the ‘Seurat’ package, we profiled the transcriptome of 19 distinct hippocampal lineages. This resolution was set to establish a balance between over-clustering and sub-population level classification of abundant lineages such as pyramidal cells and oligodendrocytes (Figure 5A). With this approach, our analysis segregates pyramidal neurons into 6 distinct layers including newly formed migratory (NFM), newly formed ventral (NFV), CA1-dorsal (CA1-D), CA1-ventral (CA1-V), CA2 and CA3 lineages. Additionally, oligodendrocyte and oligodendrocyte progenitor cells (OPC) were distinguishable into 2 lineages each. The rest of the lineages resolved here, i.e. dentate gyrus (DG), intermediate neurons, interneurons, microglia, astrocytes, pericytes, endothelial, ependymal, CD274+ neuroimmune cells, were classified as they exist in the original study (Singh et al., 2023). Finally, the high fidelity of cell type identification was set by cross-validating the pattern of observed differentially expressed genes (DEGs) with known markers (Figure 5B). Most of these DEGs were concomitant with known *bona fide* markers that are shown to be vital elements in the classified lineages (details provided upon request). Intriguingly, from the repertoire of DEGs, we find that the upregulation of *Zbtb20* was flagged in the ‘intermediate cell’ cluster. ‘Intermediate cells’ were identified based on the upregulation and co-expression of DG specific genes (*Prox1 and Rfx3*) and developing pyramidal neuronal genes (*Nfia and Rgs6*). Moreover, *Sema3c,* a gene that assists in developing the physiology of pyramidal neurons (Shane, 1976), was exclusively expressed in ‘Intermediate cells’ amongst all neuronal lineages. Of note, in consistency with genome wide *in situ* hybridization image data (mouse.brain-map.org), *Zbtb20* expression is perceivable in DG, CA1-D, CA3, astrocytes and vascular cells (Figure 6A-B). Taken together, the upregulation of *Zbtb20* marks the intermediate cells that seem to link DG and developing pyramidal neuronal populations which accords with its postulated function of neuroplasticity in health and pathologies.

**Figure 5:**
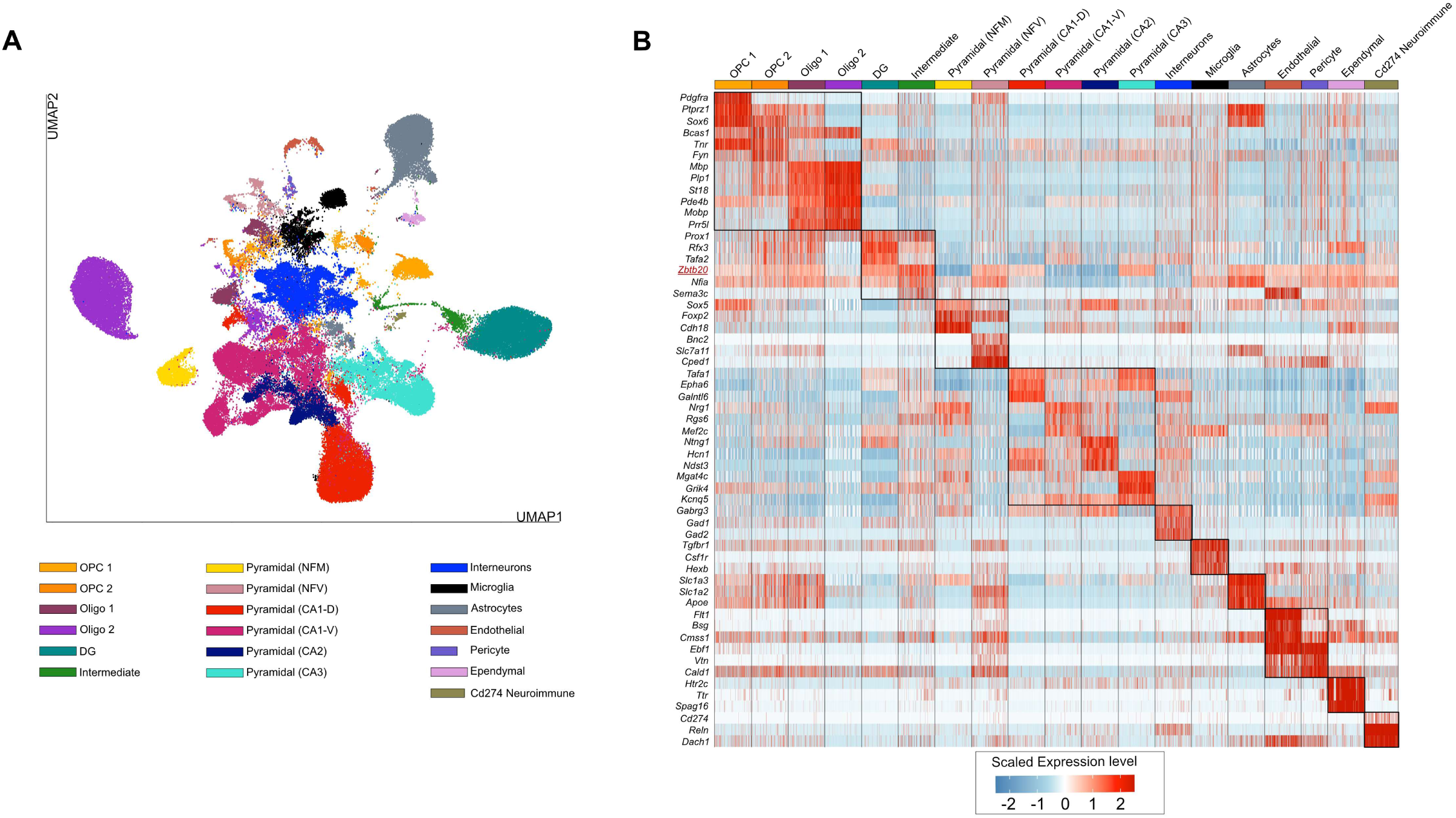
Classification and molecular markers of distinct lineages in the mouse hippocampus. **(A)** 2-dimensional Uniform Manifold Approximation Plot (UMAP) resolving the 19 distinct lineages of mouse hippocampus [integrated snRNA-seq derived from rhEPO (N=6) and placebo (N=6) subjects]. Cluster identity was assigned manually by retrieving known marker genes. The major cell types are annotated on the plot. Each dot represents a nucleus. **(B)** Heatmap illustrating the relative expression of the top 6 key genes defining the identity of each cluster in **A**. The top differentially expressed genes were calculated using Wilcoxon rank-sum test for each identified cell type. The color ranges from low (blue) to high (red) expression of marker genes (rows) in the single nuclei (columns). Full list of DEGs in each cell type are given in Supplementary Table 3.

Next, we asked whether rhEPO treatment affects the *Zbtb20* expression pattern in any or many of the aforementioned lineages. By comparing the transcriptome of identical cell types between rhEPO and placebo subjects, we identified the significant dysregulation of *Zbtb20* in multiple cell types. While the dysregulation was nominal in vascular and neuroimmune cells, pyramidal (NFM) neurons exhibited the dramatic effect of rhEPO on *Zbtb20* expression (Figure 6C). Most of pyramidal (NFM) cells expressing *Zbtb20* are from rhEPO samples (Figure 6D) indicating that *Zbtb20* expression is induced upon rhEPO specifically in 1 lineage. Because pyramidal (NFM) cells are highly specialized in combating the disorders of mood, memory and cognition, and in the potential to intricately integrate in neural circuits and the induction of immediate early genes upon metabolic challenges, our results point towards a new player (ZBTB20) for therapeutic purposes. Considering the transcription factor activities of ZBTB20, overall, we show the involvement of ZBTB20 in linking neuronal activity to gene expression and plasticity that is induced by rhEPO.

**Figure 6:**
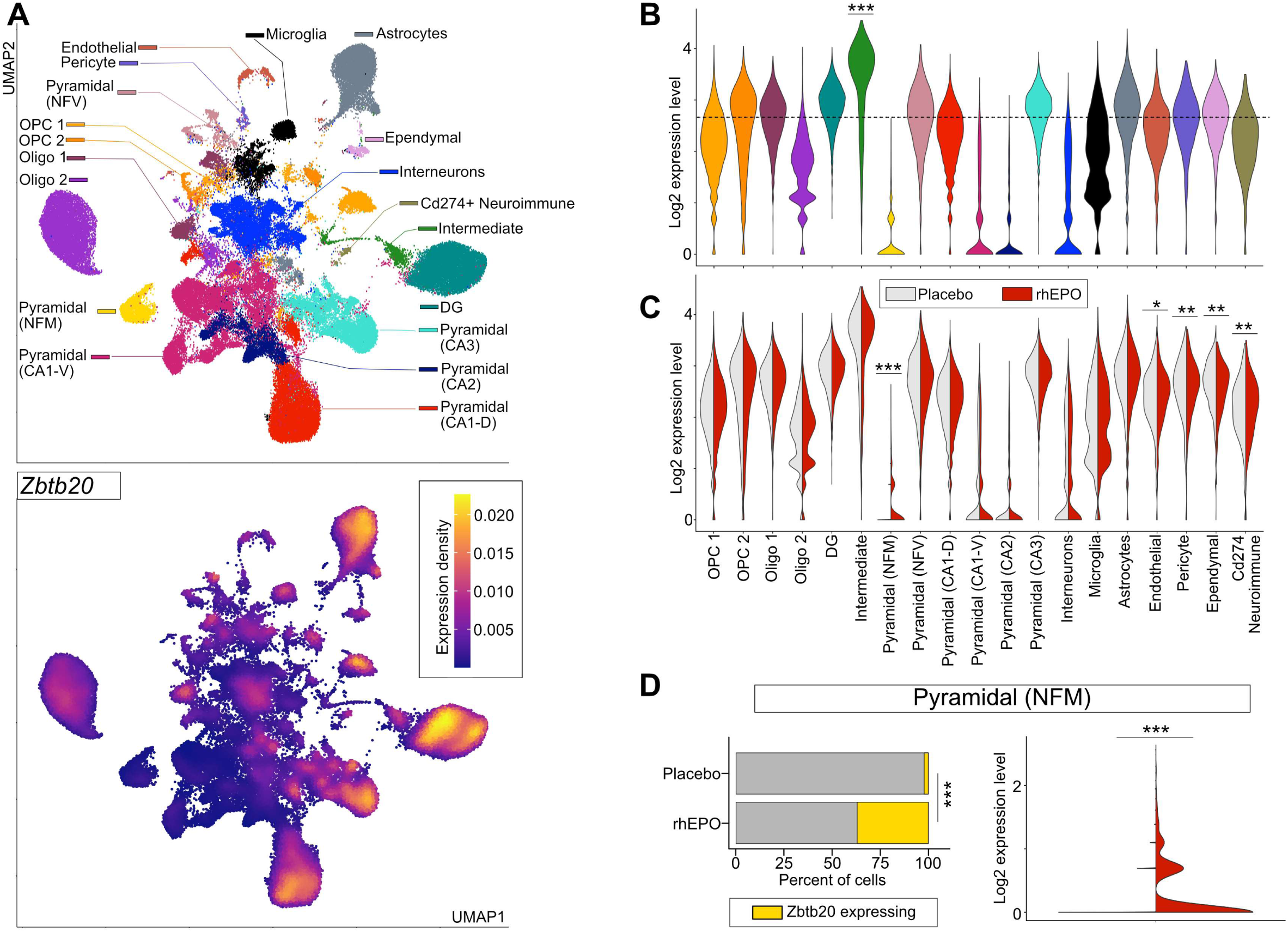
rhEPO induces the ZBTB20 expression in the newly formed migratory pyramidal lineage. **(A)** Feature plot (lower panel) based on UMAP (upper panel) demonstrating the wider pattern of ZBTB20 levels, denoted as the density of relative expression. Midnight blue area represents lower expression whereas the nuclei under the golden area exhibit higher expression of ZBTB20. Note: ZBTB20 expression in multiple premises suggesting the intricacy of ZBTB20 regulation in hippocampus needs to be unveiled in future studies. **(B)** Violin plot showing the expression distribution of ZBTB20 across the lineages coded by colors as denoted in **A**. Stars signify the adjusted p-values obtained by Benjamini-Hochberg (BH) correction followed by Wilcoxon rank-sum test (1-way), comparing ZBTB20 expression in ′Intermediate cells′ versus the rest of lineages (*** < 0.01, BH test). **(C)** Grouped violin plot visualizes the comparison of expressional dynamics of ZBTB20 between placebo (light grey) and rhEPO (red) across lineages; level of significance is calculated similarly as shown in previous plot, except, here, the comparison is made between placebo and rhEPO samples (* < 0.05; *** < 0.01, BH test). **(D)** Left panel: Stacked bar plot of pyramidal (NFM) cell type illustrating the percent of cells expressing ZBTB20 (gold) and the remaining cells (grey) separately in placebo and rhEPO samples. Star denotes the p-value of difference in cells expressing ZBTB20 between rhEPO and placebo samples (*** < 0.01, 2-sided Fischer exact test); Right panel: Violin plot quantifying the density and distribution of ZBTB20 expression in pyramidal (NFM) cell type from rhEPO and placebo samples in pairwise manner (***< 0.01, Wilcoxon rank-sum test, 1-way).

## Discussion

The objective of the present study was to provide the first comprehensive behavioral characterization of ZBTB20 deficient mice, a construct-valid animal model of Primrose syndrome, and to shed light on rhEPO as a potential therapeutic intervention. We report here that *Zbtb20^+/-^* mice exhibit primary ASD-like behavior, comprising impairments in ASD core domains sociability and communication, despite unaffected olfaction, as well as repetitive conduct (Zoghbi & Bear, 2012). Second-line symptoms associated with ASD, such as hyposensitivity towards heat-mediated nociception, compromised cognitive flexibility, and hyperactivity, are additionally observed. Notably, hyperactivity is described here for the first time in murine ZBTB20 deficiency, while impaired cognition and hyposensitivity for heat-mediated pain have previously been identified in *Zbtb20^+/-^* mice (Jia et al., 2021; Nielsen et al., 2007). Intriguingly, rhEPO treatment indeed attenuates the *Zbtb20^+/-^* phenotype, namely enhances sociability, cognitive flexibility, working memory, as well as the drive and motivation to explore.

The abundance of ZBTB20 in hippocampus (Nielsen et al., 2014) has led this and other studies of ZBTB20 deficient mice (Gulbranson et al., 2021; Ren et al., 2012) to investigate hippocampus-dependent properties which revealed impaired learning and memory performance in various behavioral tests. This was similarly found in human patients with Primrose syndrome. Here, several children and adults were also diagnosed with ASD and/or ADHD in the sense of a double-diagnosis (Melis et al., 2020; Rasmussen et al., 2014). Since prior data indicated an increase in ZBTB20 expression in hippocampal CA1 neurons following rhEPO treatment (Fernandez Garcia-Agudo et al., 2021), we hypothesized that rhEPO might attenuate the *Zbtb20^+/-^* phenotype. Indeed, sociability, cognitive flexibility, working memory, and motivation to explore improved in *Zbtb20^+/-^* mice upon rhEPO. Previously, beneficial effects of EPO on social interaction in rats were demonstrated in an ASD model (Hosny et al., 2023) and in models of perinatal brain injury (Jantzie et al., 2018; Robinson et al., 2018). Cognitive flexibility, assessed by reversal tasks in Morris Water Maze, was compromised in *Zbtb20^+/-^* males and, in agreement with other studies (Robinson et al., 2018; Robinson et al., 2022), rhEPO clearly enhanced their respective performance. Analogously, superior cognitive flexibility was observed in mice with a constitutively active EPO receptor (Sargin et al., 2011), and rhEPO markedly increases hippocampus-dependent learning and memory (Adamcio et al., 2008; Almaguer-Melian et al., 2015; Garmabi et al., 2022; Hamidi et al., 2013; Jacobs et al., 2021; Jia et al., 2016; Kumral et al., 2004; Leconte et al., 2011; Lu et al., 2005; Ma et al., 2018; Yu et al., 2015).

EPO has a promoting impact on working memory in ZBTB20 deficient mice as shown in the Y-Maze test. This aligns with a mouse study on rhEPO in traumatic brain injury (Chauhan & Gatto, 2010). Furthermore, a synthetic EPO derivative without erythropoietic effect improved working memory in a rat model of multiple sclerosis (Dmytriyeva et al., 2016). Patients with treatment-resistant major depression exhibited amended working memory upon rhEPO in an fMRI task (Miskowiak et al., 2016). Exploratory behavior, evaluated by the hole board test, delivered a surprise: *Zbtb20^+/-^* males demonstrated a lower number of rigorous revisits, an unexpected phenotype that was assuaged by EPO treatment. This could be in some line with rat studies, reporting mitigation by rhEPO of adverse effects like anxiety upon stress (Fathi et al., 2022) or upon transient middle cerebral artery occlusion (Larpthaveesarp et al., 2021). A negative correlation between EPO levels and anxiety rating was found in patients with generalized anxiety disorder (Kurutas, 2023). Attenuating the ASD phenotype in ZBTB20-deficient mice, rhEPO might be a promising treatment approach for Primrose syndrome.

The marble burying test appraises stereotypical and repetitive behaviors (Dere et al., 2014). Unexpectedly, *Zbtb20^+/-^* mice of both genders exhibited a markedly decreased number of buried marbles. Therefore, we replicated this test in a second experimental cohort and incorporated additional examinations to investigate potential reasons for this unforeseen phenotype. Once again, *Zbtb20^+/-^* mice demonstrated a decreased number of buried marbles. To rule-out anxiety-like behavior towards novel objects, the neophobia test and to exclude anhedonia and depression-like behavior, sucrose preference, tail suspension and forced swim test were performed, all revealing normal conduct. Even though just few cases of individuals with Primrose syndrome, co-diagnosed with anxiety, are known (Juven et al., 2020; Posmyk et al., 2011; Shuvarikov et al., 2013), future studies might want to include tests for anxiety-like behavior in mice. This needs careful planning as these tests require them to be naïve to handling (Lapin, 1995).

The present study employed for the first time MRI volumetric analyses of the brain in an animal model of *Zbtb20* deficiency. It uncovered increased volumes of hippocampus, cerebellum, brain matter, and whole brain in ZBTB20 deficient males. Weight assessment of *Zbtb20^+/-^* male brains immediately after extraction and following lyophilization showed an increase in both weight and dry weight. In contrast to *Zbtb20^+/-^* mice, MRI scans of human subjects diagnosed with Primrose syndrome predominantly exhibited malformation or agenesis of corpus callosum. There were also sporadic cases of colpocephaly, Chiari malformation, and partial calcification of the basal ganglia (Arora et al., 1993; Cleaver et al., 2019; Dalal et al., 2010; Long et al., 2023). While most studies of Primrose syndrome report on macrocephaly (Alby et al., 2018; Arora et al., 1993; Carvalho & Speck-Martins, 2011; Casertano et al., 2017; Cleaver et al., 2019; Ferreira et al., 2019; Grimsdottir et al., 2019; Juven et al., 2020; Mathijssen et al., 2006; Melis et al., 2020; Shuvarikov et al., 2013; Stellacci et al., 2018; Yamamoto-Shimojima et al., 2020), only 2 describe megalencephaly (Abdallah et al., 2024), which aligns with the findings of the present study. The observed imbalance between reports of macrocephaly and megalencephaly might be attributed to the different methods applied. While macrocephaly can be determined through a simple measurement of head circumference, diagnosing megalencephaly requires either *in vivo* imaging or post mortem tissue analyses. Previously, a reduction in wet and dry weight of lungs and hearts of *Zbtb20^-/-^* mice was reported (Ren et al., 2020) which, however, disappeared when normalized to body weight or tibia length. While *Zbtb20^+/-^* males exhibited an increase in body weight and tibia length, there were no differences in weight of lungs and hearts. Obviously, the genotype is the driving factor, with homozygous ZBTB20 deficiency causing a more severe impact on lungs and hearts.

While *Zbtb20^+/-^* males displayed a pronounced ASD phenotype, females did just partially. Among the social tests conducted, only nest building revealed abnormalities, showing reduced quality of nests in both placebo and rhEPO-treated *Zbtb20^+/-^* groups. Similar to males, females demonstrated repetitive behavior due to excessive climbing in their home cages and hyposensitivity towards heat-mediated nociception, the latter being rescued by rhEPO treatment. While *Zbtb20^+/-^* males exhibited impairments in spatial learning and memory, as well as reversal learning and memory in the Morris water maze, *Zbtb20^+/-^* females showed only difficulties with spatial and reversal memory, in agreement with *Zbtb20^+/-^* females in the Barnes maze (Nielsen et al., 2007). A lack of motivation to explore in females was also evident in the tendency towards a reduced number of rigorous revisits and marbles buried, as was the phenotype of hyperactivity. Unlike in males, thermal analysis during the 3-chamber sociability test revealed a decreased centralization index during sociability testing, with no differences observed during habituation. This suggests a reduced stress level in *Zbtb20^+/-^* females when forced into a situation of social interaction. MRI revealed a reduced olfactory bulb volume in both placebo and rhEPO-treated *Zbtb20^+/-^* females. The volumes of hippocampus and cerebellum were higher in placebo-treated *Zbtb20^+/-^* females and reduced to control levels under rhEPO conditions. Additionally, ventricle volumes were reduced in rhEPO-treated mice.

In human studies, 50% of individuals diagnosed with Primrose syndrome were reported to be male, 40% female, and 10% without stated sex (Abdallah et al., 2024; Alby et al., 2018; Arora et al., 2020; Battisti et al., 2002; Carvalho & Speck-Martins, 2011; Casertano et al., 2017; Cleaver et al., 2019; Dalal et al., 2010; Ferreira et al., 2019; Grimsdottir et al., 2019; Juven et al., 2020; Lindor et al., 1996; Mathijssen et al., 2006; Mattioli et al., 2016; Melis et al., 2020; Posmyk et al., 2011; Rasmussen et al., 2014; Shuvarikov et al., 2013; Stellacci et al., 2018; Yamamoto-Shimojima et al., 2020), indicating a sex-independent distribution of Primrose syndrome across the population.

## Conclusions

The present study offers for the first time an in-depth behavioral characterization of ZBTB20 deficient mice, which exhibit ASD-like behavior. MRI revealed structural changes in specific brain regions. Together, these results suggest ZBTB20 deficient mice as a valuable experimental model for Primrose syndrome. Importantly, 3 weeks of rhEPO treatment, initiated at PND28, ameliorates the phenotype. This puts rhEPO in the spotlight as potential treatment for Primrose syndrome.

## Acknowledgements

This work has been funded by the European Research Council (ERC) Advanced Grant to HE under the European Union’s Horizon Europe research and innovation programme (acronym *BREPOCI;* grant agreement No 101054369). Furthermore, the study has been fostered by the Max Planck Society and the Max Planck Förderstiftung. Research of HE is supported by DFG TRR-274/1 2020-408885537. VDG received backing from the IMPRS-Genome Science PhD program. The authors thank Lars van Werven for his assistance in tissue lyophilization and Dr. Jan Seidel for providing the original scripts to analyze images acquired by thermal cameras. Research of ABT is supported by NextGenerationEU via Bulgarian National Recovery and Resilience Plan, Project #BG-RRP-2.004-0009-C03 (MUVE-TEAM).

## Author Contributions

Supervision: HE

Conception and design: MH, HE

Funding acquisition: HE, KAN

Data acquisition/analysis/interpretation: MH, AR, VG, JW, YC, UB, SO, KH, MS, RD, UÇ, SB, AS, ABT, KAN, HE

Drafting the manuscript: MH, HE Drafting display items: MH, JW, MS, HE

All authors read and approved the final version of the paper.

## Data Availability Statement

All reported data are available upon informed request.

## Abbreviations

ACK: Ammonium-Chloride-Potassium
ADHD: attention deficit/hyperactivity disorder
ASD: autism spectrum disorder
BW: body weight
CA1: cornu ammonis 1
CI: centralization index
DG: dentate gyrus
DPI: days post injection
FACS: fluorescent-activated cell sorting
IHC: immunohistochemistry
LABORAS: laboratory animal behavior observation registration and analysis system
MRI: magnetic resonance imaging
NFM: newly formed migratory
PBS: phosphate-buffered saline
PND: postnatal day
rhEPO: recombinant human erythropoietin
ROIs: regions of interest
RRR: replacement-reduction-refinement
RT: room temperature
snRNA-seq: single nuclei RNA-sequencing
WT: wild type
ZBTB20: zinc finger and BTB domain-containing protein 20.

---

**Supplementary Figure 1:**
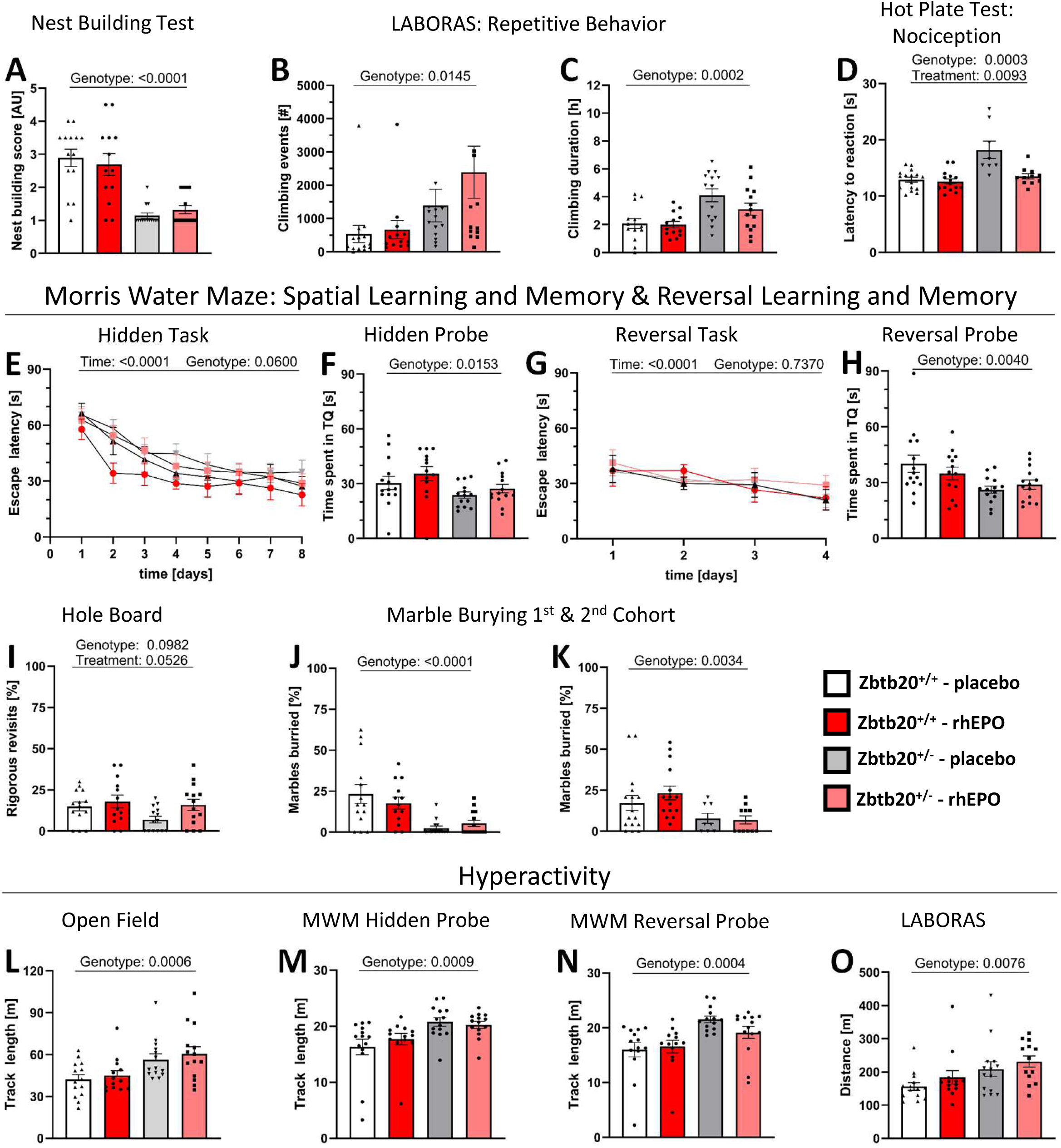
Behavioral phenotype of *Zbtb20^+/-^* females. **(A)** Nest building, a social test, shows reduced scores in placebo- and rhEPO-treated *Zbtb20^+/-^*^-^ females. **(B-C)** Repetitive behavior is demonstrated in *Zbtb20^+/-^* females in LABORAS via increased events **B** and duration **C** of climbing. **(D)** Hot plate test reveals hyposensitivity for heat-mediated nociception in placebo-treated *Zbtb20^+/-^* females, while rhEPO-treated *Zbtb20^+/-^* mice demonstrate behavior on control level. **(E-H)** Assessment of spatial learning and memory in Morris Water Maze. While 8 days of hidden task training show no genotype-based differences **E**, hidden probe trial divulges reduced time spent in target quadrant (TQ) for both *Zbtb20^+/-^* groups **F**. Subsequent 4 days of reversal training demonstrate no genotype effect across groups **G**, while reversal probe trial discloses reduced time spent in TQ for both *Zbtb20^+/-^* groups **H**. **(I)** Placebo-treated *Zbtb20^+/-^* females demonstrate reduced number of rigorous revisits in hole board testing, while rhEPO-treated *Zbtb20^+/-^* females express behavior on control level. **(J-K)** Both *Zbtb20^+/-^* female groups show reduced number of marbles buried **J,** an unexpected result, confirmed **K** in an additional cohort. **(L-O)** Increased activity of both placebo- and rhEPO-treated *Zbtb20^+/-^* females is observable in the track lengths of open field (**L**), both probe trials of Morris water maze (**M, N**), as well as LABORAS (**O**); (2-way or 3-way ANOVA as applicable; mean±SEM).

**Supplementary Figure 2:**
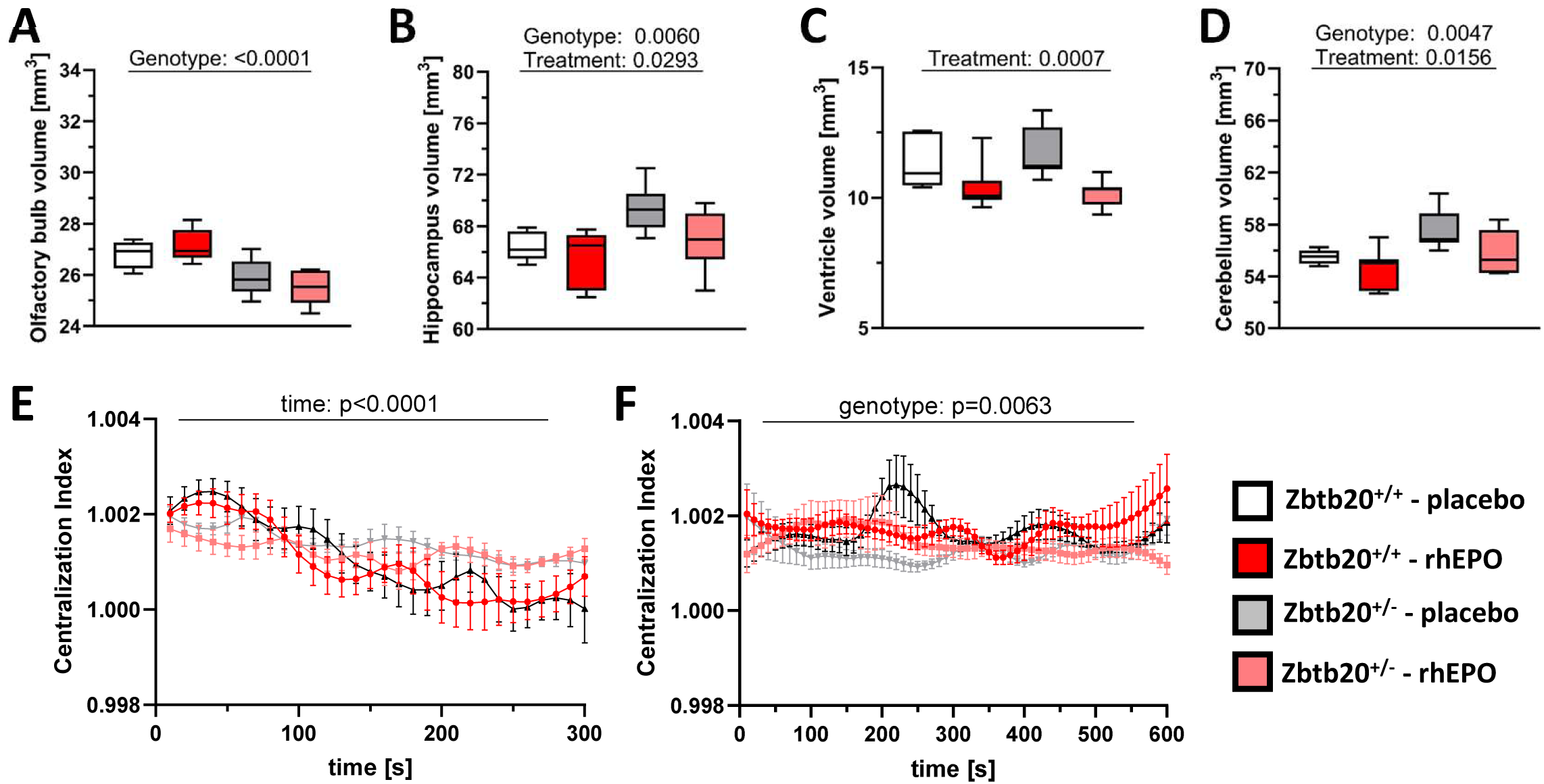
Behavior of *Zbtb20^+/-^* females – continued. **(A-D)** MRI reveals structural differences in brain areas of *Zbtb20^+/-^* females, such as olfactory bulb **A**, hippocampus **B**, ventricles **C**, and cerebellum **D**. **(E-F)** Thermal-based analysis during 3-chamber sociability test discloses only a time effect of the centralization index during habituation **E**, but a reduced centralization index during trial of both *Zbtb20^+/-^* groups **F**; (2-way or 3-way ANOVA as applicable; mean±SEM).

**Supplementary Figure 3:**
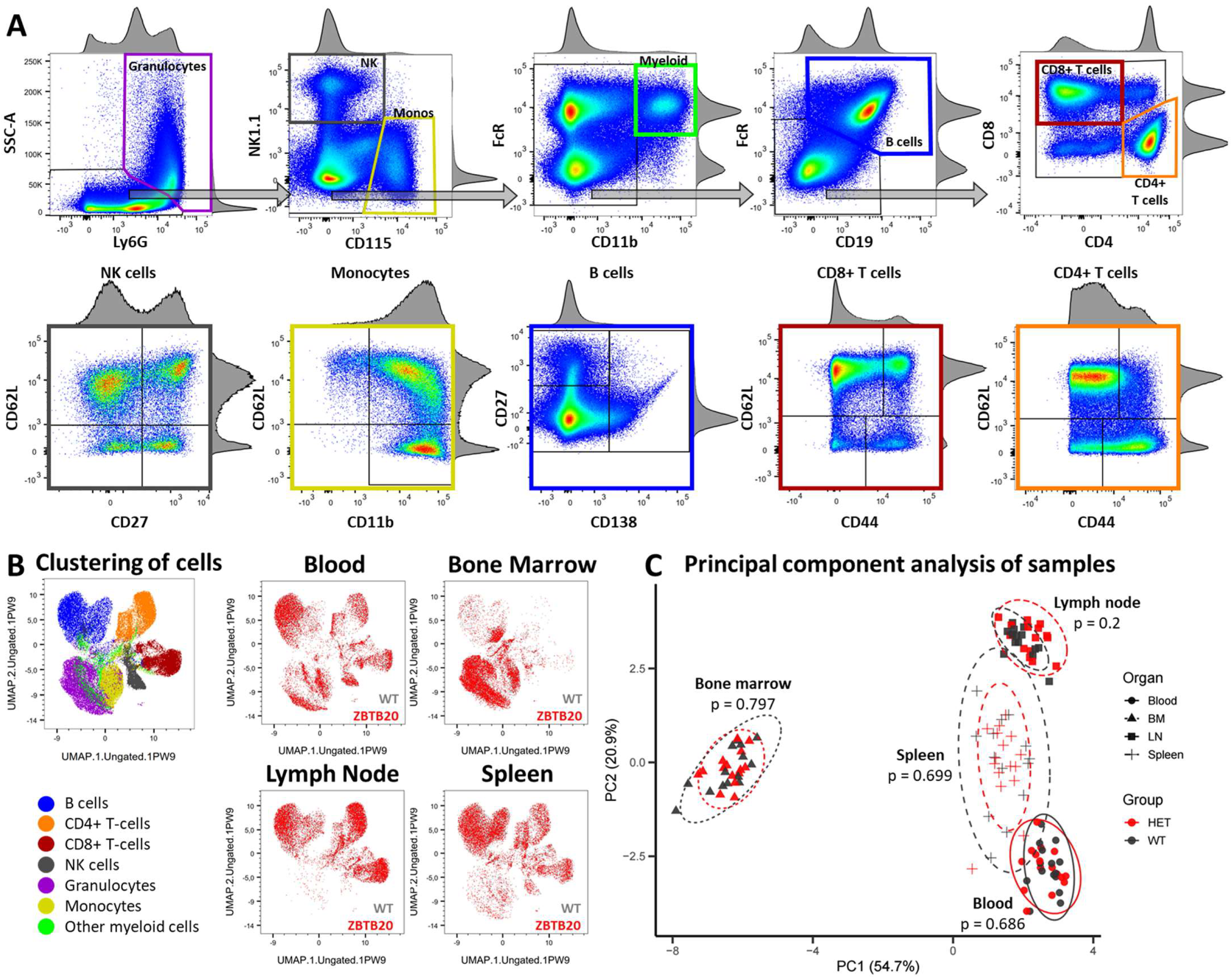
Immunophenotyping of major lymphoid compartments. **(A)** Gating strategy for the determination of 6 major (color coded) and 19 minor immune cell subsets. Arrows indicate the hierarchy of gating. **(B)** UMAP representation of the 6 major immune cell subsets and 1 minor immune cell subset and qualitative comparison of immunophenotypes of *Zbtb20* haploinsufficient (ZBTB20, red) and wildtype (WT, gray) mice in blood, bone marrow, lymph node, and spleen. While the 4 lymphoid compartments show distinct distributions of white blood cells, haploinsufficient ZBTB20 mice display no gross abnormalities in comparison to wildtype mice. All graphs show the concatenated data generated by concatenating 10000 live hematopoietic single cells per sample (N=120, n=14-16 per genotype and lymphoid organ). **(C)** Principal component analysis of z-scored immune cell frequencies. Each point represents one mouse within a given organ; 95% confidence ellipses are shown for each genotype and organ combination. Global differences in immune composition between *Zbtb20^+/-^* and WT mice were tested separately for each organ using PERMANOVA on Euclidean distances of the scaled data (p values shown).

**Supplementary Tables 1 & 2**

**Complete overview of all parameters on behavior & MRI - males & females (1) Detailed data on *Zbtb20^+/-^* males.** Behavioral analyses, MRI, organ weights and qPCR presented for all 4 groups across the project [mean, standard deviation (SD), group sizes (n)]. Applied statistical tests were t test with welch correction (t), Mann-Whitney-U test (U), and 1-, 2-, or 3-way-ANOVA. P values below the determined significance level of 0.05 are indicated in bold.

**(2) Detailed data on *Zbtb20^+/-^* females.** Behavioral analyses, MRI, organ weights and qPCR presented for all 4 groups across the project [mean, standard deviation (SD), group sizes (n)]. Applied statistical tests were t test with welch correction (t), Mann-Whitney-U test (U), and 1-, 2-, or 3-way-ANOVA. P values below the determined significance level of 0.05 are indicated in bold.

**Supplementary Table 3:**

**Full list of differentially expressed genes (DEGs)** with known *bona fide* markers that are shown to be vital elements in the classified cell lineages (details provided upon request).

**Supplementary Table 4:**

**High Parameter Flow Cytometry data:** Frequencies of 6 major and 19 minor immune cell subsets across 4 lymphoid compartments in wildtype (WT) versus *Zbtb20^+/-^* (heterozygous knockout; Het) mice. Data displayed in percentage and as mean ± standard deviation. Statistical testing was based on data normality and variance; t-test = two-sided t-test; Wcc = Wilcoxon rank sum test with continuity correction; Welch = Welch’s corrected two sample t-test. FDR = false discovery rate. d = Cohens’ d. N=14-16 mice per genotype.

